# Housekeeping gene *gyrA*, a potential molecular marker for *Bacillus* ecology study

**DOI:** 10.1101/2022.04.11.487959

**Authors:** Yan Liu, Polonca Štefanič, Youzhi Miao, Yansheng Xue, Weibing Xun, Qirong Shen, Zhihui Xu, Ruifu Zhang, Ines Mandic-Mulec

## Abstract

*Bacillus* is a ubiquitous microorganism and is one of the most commercially important species widely used in industry, agriculture and healthcare. *Bacillus* is relatively well understood at the single-cell level; however, molecular tools that determine diversity and ecology of *Bacillus* community are limited, which limits our understanding of how the *Bacillus* community works. In the present study, we investigated the potential of the housekeeping gene *gyrA* as a molecular marker for determining the diversity of *Bacillus* species. The amplification efficiency for *Bacillus* species diversity could be greatly improved by primer design. Therefore, we designed a novel primer pair *gyrA*3 that can detect at least 92 *Bacillus* species and related species. For *Bacillus amyloliquefaciens, Bacillus pumilus*, and *Bacillus megaterium*, we observed that the high variability of the *gyrA* gene allows for more detailed clustering at the subspecies level that cannot be achieved by the 16S rRNA gene. Since *gyrA* provides better phylogenetic resolution than 16S rRNA and informs on the diversity of the *Bacillus* community, we propose that *gyrA* gene may have broad application prospects in the study of *Bacillus* ecology.

## Introduction

Microbial communities in soil are known to be one of the largest reservoirs of biological diversity and have been extensively studied (Timmis & Ramos, 2021). Current advances in high-throughput DNA sequencing of a portion of the small subunit of ribosomal RNA (16S and 18S rRNA) form the backbone of most studies of soil microbial ecology (Klindworth et al., 2013). For bacteria, the 16S rRNA gene is usually preferred, because it contains both highly conserved and hypervariable regions (Peer, Chapelle, & Wachter, 1996), and especially because comprehensive reference databases have been compiled for comparison (Cole et al., 2014; McDonald et al., 2012; Quast et al., 2013; Yoon et al., 2017). However, 16S rRNA amplicon sequencing also has many shortcomings: first, 16S rRNA evolves slowly and is highly conserved, making it a poor marker for distinguishing between closely related strains. Second, chimera formation during PCR is high because 16S rRNA variability is very low (Pinto & Raskin, 2012; Sun, Jiang, Wu, & Zhou, 2013). Third, the number of 16S rRNA copies in different species is highly variable, and single nucleotide polymorphisms (SNPs) at the single-cell level may result in an overestimation of diversity (Johnson et al., 2019). Fourth, the similarity between species can be very high, making it difficult to delineate species in cluster analysis, and different clustering levels lead to different results (Eren et al., 2013). Therefore, new complementary taxonomic markers for genetic and bioinformatic analysis need to be developed to study microbial diversity in more detail, especially at the subspecies level.

*Bacillus* is one of the most intensely studied bacterial genus comprising at least 200 species (Mandic-Mulec, Stefanic, & van Elsas, 2016). It is a heterogeneous bacterial taxon that is ubiquitous in various ecological niches and widely used in medicine, industry and agriculture. Although there have been many in-depth studies on *Bacillus* model species, community-level studies on *Bacillus* in soil and other habitats lag behind. Because sequence of 16S rRNA within *Bacillus* species is often similar, the definition and delineation of bacterial species based on the 16S rRNA gene comparison among related species in the genus *Bacillus* is unclear. Therefore, identification and typing of *Bacillus* isolates based on the 16S rRNA gene alone cannot provide accurate results and it is important to explore and use other genes as molecular markers to assess the diversity of the *Bacillus* community (Mandic-Mulec et al., 2016).

Housekeeping genes are potential candidates for assessing microbial diversity because they have been shown to elicit higher phylogenetic resolution than the 16S rRNA gene, such as the *rpoB* gene, *gyrA* gene, and *gyrB* gene, etc.(Chun & Bae, 2000; Hurtle et al., 2004; Kasai, Tamura, & Harayama, 2000; Levican, Collado, & Figueras, 2013; Ménard, Buissonnière, Prouzet-Mauléon, Sifré, & Mégraud, 2016; Stefanic et al., 2012; Stefanic, Kraigher, Lyons, Kolter, & Mandic-Mulec, 2015; Stefanic & Mandic-Mulec, 2009; Yamamoto et al., 2000). Typically, there are only one or two copies of housekeeping genes per genome, and the use of low-copy number genes compared to high-copy number genes could lead to more accurate diversity analysis by avoiding overestimation of diversity due to SNPs in different gene copies. Indeed, several studies have tested the *rpoB* gene and the *gyrB* gene as molecular markers to analyze the diversity of bacterial communities by amplicon sequencing. The results showed that housekeeping gene sequencing provided a more accurate description of bacterial community composition than 16S rRNA sequencing under certain conditions (Ogier, Pagès, Galan, Barret, & Gaudriault, 2019; Poirier et al., 2018; Vos, Quince, Pijl, de Hollander, & Kowalchuk, 2012).

The housekeeping gene *gyrA*, encoding DNA gyrase subunit A, is essential for DNA replication and present in all bacteria (Cozzarelli, 1980). Analyses of *gyrA* identity provided higher phylogenetic resolution than the 16S rRNA gene for tested *Bacillus* isolates (Chun & Bae, 2000; Ménard et al., 2016). Specifically, partial *gyrA* gene sequences was used for phylogenetic analysis and species identification of seven *Bacillus* strains, including *B. amyloliquefaciens, B. atrophaeus, B. licheniformis, B. mojavensis, B. subtilis, B. subtilis subsp. Spizizenii*, and *B. vallismortis* (Chun & Bae, 2000). Moreover, the *gyrA* gene sequences provided a good marker for *B. subtilis* and *B. amyloliquefaciens* and showed better discriminatory potential between these two closely related species than the *rpoB* gene (Chun & Bae, 2000). The *gyrA* gene has been also successfully used to detect intraspecific diversity of *B. subtilis* isolates from soil microscale (Stefanic and Mandic Mulec, 2009) and tomato rhizosphere isolates (Oslizlo et al., 2015). Overall, these works suggest that *gyrA* has a good potential to be used as a molecular marker for microbial ecology studies of *Bacillus* genus and related species, however, to the best of our knowledge, the available primer pairs used in studies indicated above, have not been used for amplicon based community analyses of *Bacillus* species.

In this study, we first compared the rate of variation of the16S rRNA and *gyrA* genes between 20 *Bacillus* species and then designed a new primer pair that specifically target *gyrA* (*gyrA*3). We then evaluate their performance by using *in silico* PCR, testing their efficiency on *Bacillus* isolates and performed SNP analysis of 16S rRNA and *gyrA* genes for selected species that were available in the NCBI database. Finally, verified *gyrA*3 primers to differentiate species and strains of *Bacillus* mock community, and compared the obtained results with those targeting 16S rRNA. Our results suggest that the *gyrA* gene is a useful molecular marker for identification of *Bacillus* isolates and describing the diversity of the *Bacillus* community.

## Results

### The *gyrA* gene of the *Bacillus* genus shows higher variation rates than 16S rRNA

The housekeeping gene *gyrA* is considered to be more variable than 16S rRNA and has been used as a molecular tool for classification and identification of *Bacillus subtilis* species (Borshchevskaya, Kalinina, & Sineokii, 2013; Chun & Bae, 2000). To determine the specific variation divergence of 16S rRNA and *gyrA* gene sequences in the genus *Bacillus*, two alignments with sequences of the entire 16S rRNA and *gyrA* genes from 20 *Bacillus* species (361 genomes) (Table S1) were used to analyze nucleotide diversity. The nucleotide diversity (Pi) of 16S rRNA and *gyrA* gene calculated by the DnaSP software was 0,039 and 0.491, respectively (Fig. 1A,B blue line). The results showed that the degree of interspecies variation was significantly higher for the *gyrA* gene than for the 16S rRNA gene.

**Figure 1.**
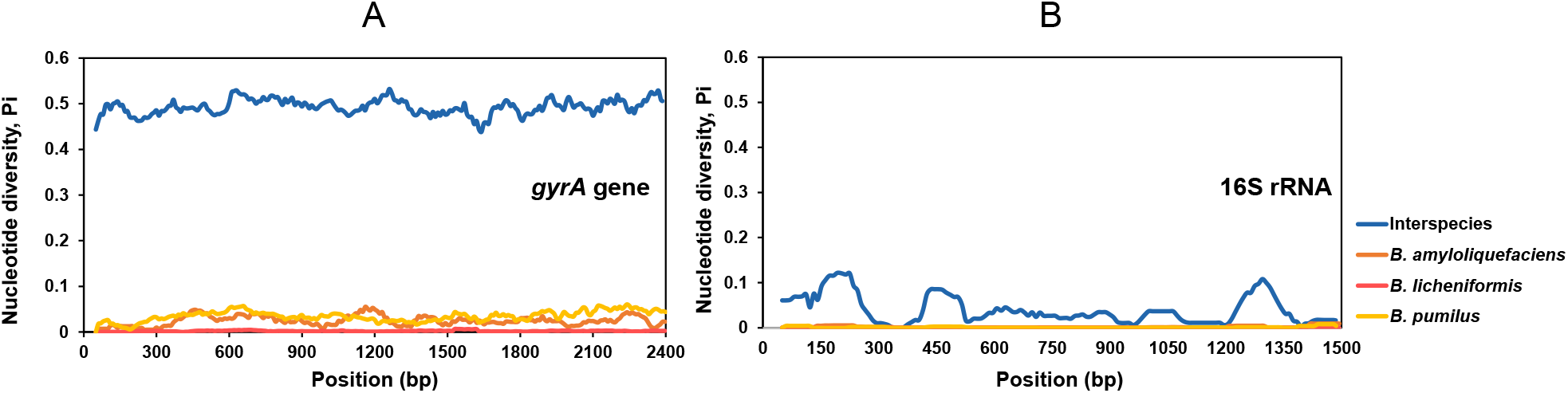
Sliding window analysis of nucleotide variability [Pi (π)] along the sequence of the *gyrA* gene (**A)** and 16S rRNA (**B)** in *Bacillus*. 361 and 256 of *gyrA* and 16S sequences from 20 species and three species (*B. amyloliquefaciens* (n=88), *B. licheniformis* (n=71) and *B. pumilus* (n=97)) were aligned with Kalign method and nucleotide diversity was determined in DnaSP (v.6) (Rozas et al., 2017) using a window size of 100-bp, a step size of 10-bp, and points based on the mid-point of each window (i.e., the first point is at position 200). The blue lines display the *Bacillus* inter-species nucleotide diversity of the two genes and the orange, grey and yellow lines display the nucleotide diversity of two genes in *B. amyloliquefaciens, B. licheniformis and B. pumilus*, respectively. The sequence length of *gyrA* gene for analysis was 2450 bp, and that of 16S rRNA was 1540bp.

Intraspecific variation analysis of the two genes was performed using the same approach but focusing on three *Bacillus* species: *Bacillus amyloliquefaciens* (88 genomes), *Bacillus licheniformis* (71 genomes) and *Bacillus pumilus* (97 genomes) (Table S1). The intraspecific nucleotide diversity (Pi) of 16S rRNA was again lower than the intraspecific nucleotide diversity of *gyrA* gene sequences in all three species: *B. amyloliquefaciens* (Pi_16S_*=*0.0014; Pi_*gyrA*_=0.0244), *B. licheniformis* (Pi_16S_*=*0.00024; Pi_*gyrA*_=0.0021) and *B. pumilus* (Pi_16S_*=*0.00136; Pi_*gyrA*_=0.0344) (Fig. 1A,B nonblue lines). In *B. amyloliquefaciens, B. licheniformis* and *B. pumilus*, the degree of variation of the *gyrA* gene was significantly higher than that of 16S rRNA. In conclusion, the *Bacillus gyrA* gene shows higher variation rates than 16S rRNA, hence we propose that *gyrA* represents a promising molecular marker for analyses of *Bacillus* community diversity analyses and the diversity of *Bacillus* isolates.

### First comparative tests of three primer pairs for the detection of *Bacillus* species

As indicated above *Bacillus* isolates have been already analyzed by primers targeting *gyrA*, however the specificity of these primers has not been investigated broadly (Ansaldi, Marolt, Stebe, Mandic-Mulec, & Dubnau, 2002; Roberts, Nakamura, & Cohan, 1994). In order to satisfy the amplicon sequencing requirements, we designed a new primer pair (*gyrA*3) (Fig. 2A), and compared its amplification potential in colony PCR and virtual PCR with the previously designed primers, referred to here as *gyrA*1 (Stefanic & Mandic-Mulec, 2009) and *gyrA*2 (De Clerck et al., 2004).

**Figure 2.**
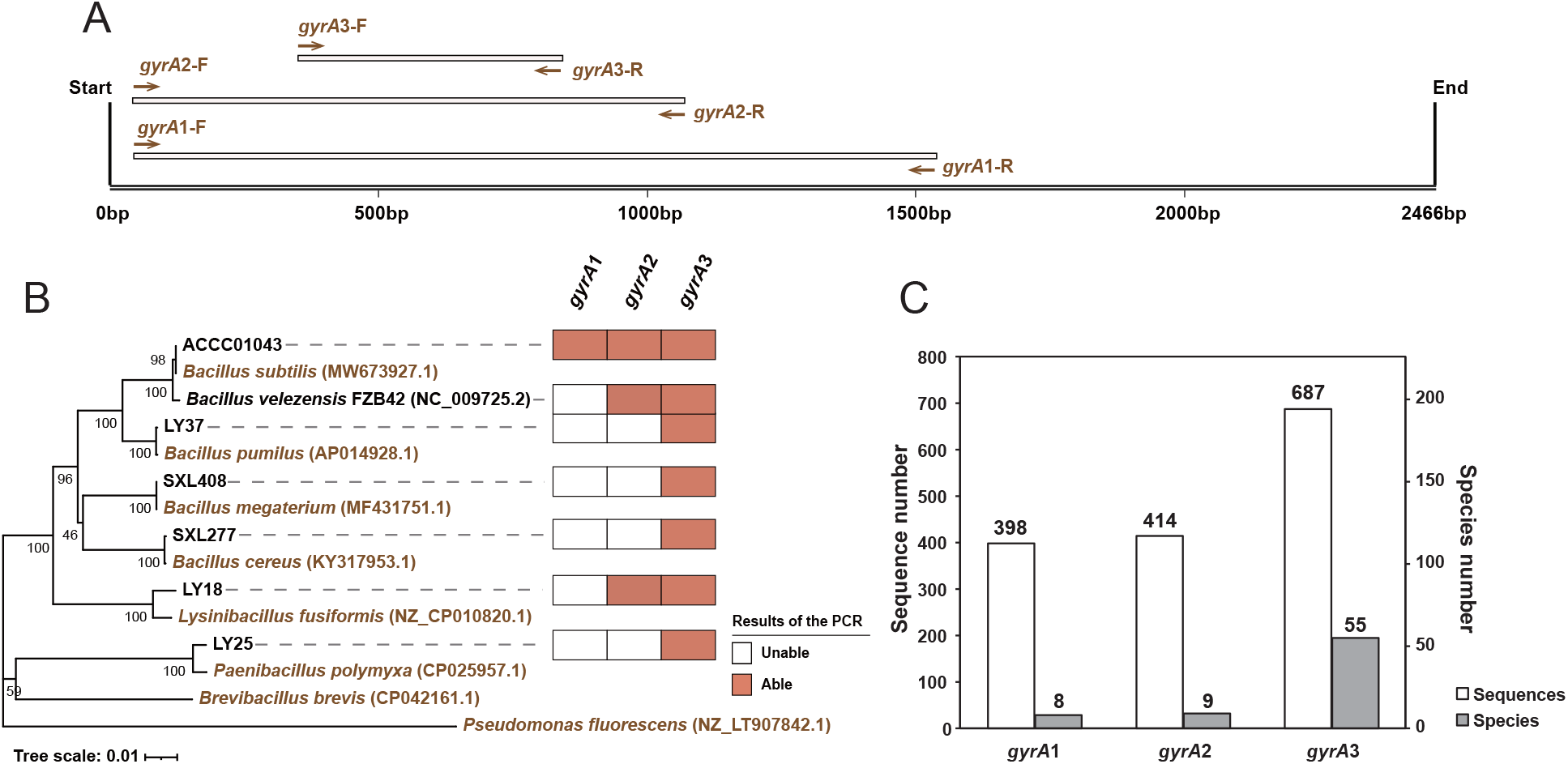
Display of the ability of the three primer pairs to amplify *gyrA* gene for seven *Bacillus* strains. (**A)** The position of *gyrA* amplicons obtained by primer pairs *gyrA*1, *gyrA*2 and *gyrA*3. (**B)** PCR amplification of seven *Bacillus* strains using three *gyrA* gene primer pairs. The orange square indicates that the target gene of *gyrA* was amplified by the indicated primer pair, and the white square indicates that it was not. The phylogenetic tree was constructed based on the complete 16S rRNA (1358bp) (Dataset S1) by using Neighbor-Joining method in MEGA 5.05 software. The reliability of clades was tested by the 1000 bootstrap replications. (**B)** Display of the results of the computer simulation amplification ability of the three *gyrA* gene primer pairs using the *Bacillus* database. The white columns represent number of sequences in the database that were matched by the primers, while the grey columns represent number of *Bacillus* species identified by the primer pair (Table S3). The database sequences were downloaded from NCBI genome database (Table S2).

First, we selected seven strains of different *Bacillus* species: *Lysinibacillus fusiformis, Paenibacillus polymyxa, Bacillus pumilus, Bacillus velezensis, Bacillus megaterium, Bacillus cereus* and *Bacillus subtilis* (Fig. 2B) to perform PCR amplification with primer pairs *gyrA*1, *gyrA*2 and *gyrA*3. The PCR amplification results showed that *gyrA*1 detected only *B. subtilis*; *gyrA*2 detected *B. subtilis, B. velezensis* and *L. fusiformis*; whereas *gyrA*3 performed much better and detected all *Bacillus* species included in the analysis (Fig. 2B and Fig. S1).

For *in-silico* PCR analysis, we constructed a *gyrA* gene library containing 5062 full-length *gyrA* gene sequences from 226 *Bacillus* species obtained from the National Center for Biotechnology Information (NCBI) (Table S2). Our criteria for positive *in-silico gyrA* PCR amplification were annealing of the primer pair with at least 18 bases match (forward primer and reverse primer). The results showed that only 8 *Bacillus* species were amplified in-silico by *gyrA*1, 9 *Bacillus* species were amplified by *gyrA*2 (Fig. S2), while 55 *Bacillus* species were amplified by *gyrA*3 (Fig. 2C). The majority of sequences amplified by *gyrA*1 and *gyrA*2 belonged to *B. subtilis*, whereas *gyrA*3 demonstrated broader diversity as evidenced by the amplification of seven species and *in-silico* PCR (Fig. 2B,C and Table S3).

### Specificity range of the *gyrA*3 primer pair by using PCR and *in silico* PCR

Because the *gyrA*3 primer pair performed better than the previously reported primers, we next combined analysis of the *in-silico* amplified *gyrA* genes with PCR analysis of *Bacillus* isolates from our laboratory culture collection. Virtual *gyrA*3 PCR amplicons from 55 different *Bacillus* species from the NCBI database were evenly distributed among the branches of the phylogenetic tree (Fig. 3 orange and green).

**Figure 3.**
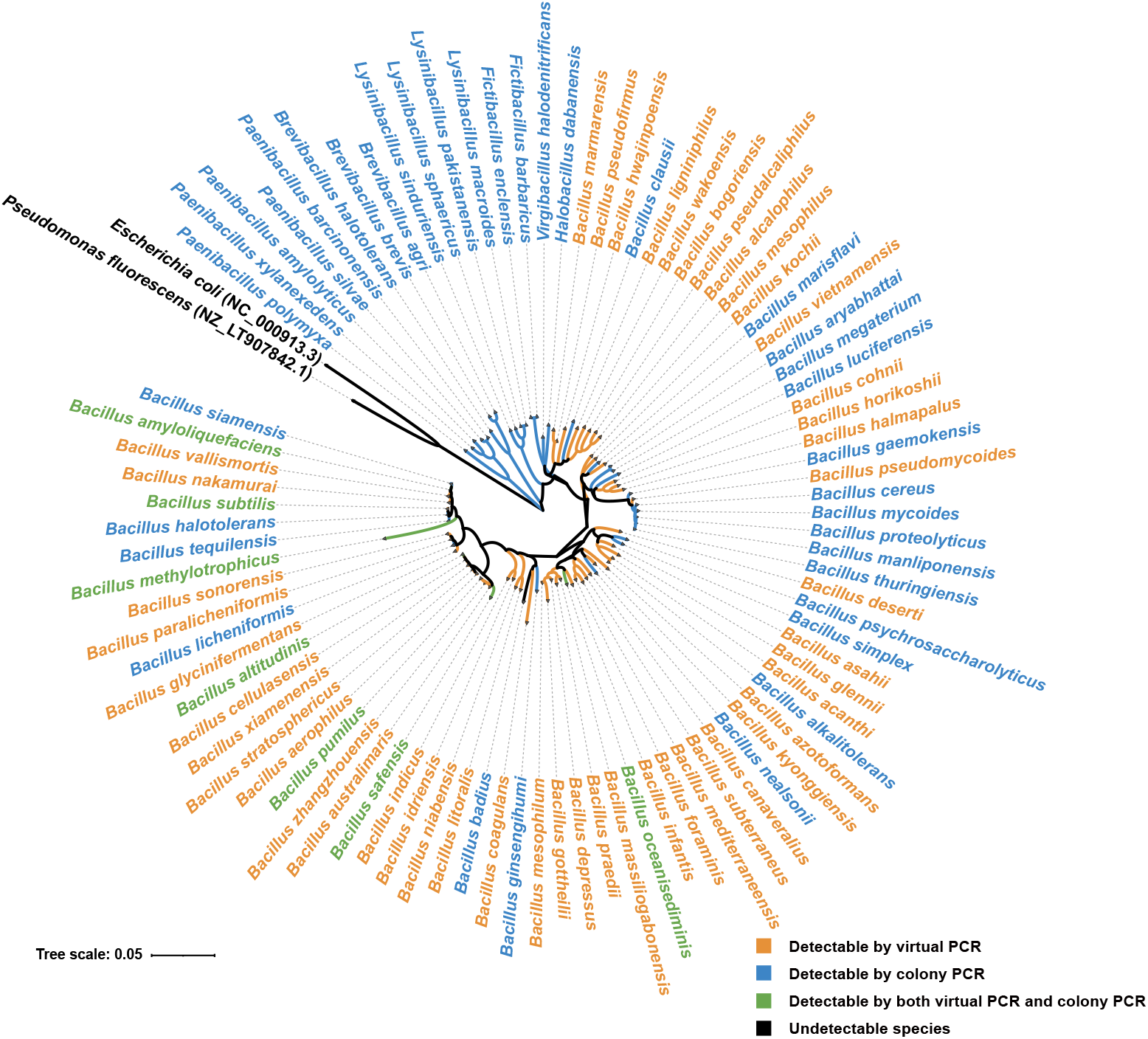
The amplification ability of the primer pair *gyrA*3 of *Bacillus* species and related species performed by computer simulation and *in situ* amplification. The *Bacillus* species indicated by the orange color (48 species) represent the species in the database that were matched by *gyrA*3 only in virtual PCR. The *Bacillus* species in blue color represented strains that can be amplified by *gyrA*3 only by colony PCR. These strains included 21 *Bacillus* species and 16 *Bacillus*’s related genera species. The *Bacillus* species in green color (7 species) represent strains that can be amplified by two methods: virtual PCR and colony PCR. The *Bacillus* species in black color (2 species) represent strains that cannot be amplified in virtual PCR nor by colony PCR. The phylogenetic tree was constructed based on 16S rRNA gene (1302bp).

Next, we used 127 *Bacillus* strains and related genera (*Paenibacillus, Lysinibacillus, Aneurinibacillus, Virgibacillus, Brevibacillus, Halobacillus, Fictibacillus*) from our laboratory culture collection to amplify their *gyrA* genes with a *gyrA*3 primer pair (Table S4). The results showed that 28 *Bacillus* species and 16 *Bacillus*-related species could be amplified by the *gyrA*3 primers (Fig. 3 blue and green). These *Bacillus* species, marked in green in the phylogenetic tree, 7 species were detectable by both methods (Fig. 3 green). In summary, the primer pair *gyrA*3 can potentially detect 76 *Bacillus* species and as many as 16 species from related genera (Fig. 3).

### The *gyrA* gene provides better intraspecific phylogenetic resolution than the 16S rRNA gene among certain species

Compared to 16S rRNA, the molecular evolution rate of *gyrA* gene sequences is faster (Timmis & Ramos, 2021), so we hypothesized that *gyrA* might provide better phylogenetic resolution at the subspecies level. To analyze the intraspecific variability of the 16S rRNA and the *gyrA* gene, we downloaded the genomes of four *Bacillus* species, each of which had more than 100 genomes available in the NCBI database, including *B. amyloliquefaciens* (116 genomes), *B. pumilus* (140 genomes), *B. megaterium* (117 genomes) and *B. anthracis* (226 genomes) (Table S5). We did not include analysis the of *B. subtilis* genomes because templates for *gyrA* primers have already been developed and applied for analyses of this species (Ansaldi et al., 2002; De Clerck et al., 2004; Roberts et al., 1994; Stefanic & Mandic-Mulec, 2009).

In *B. amyloliquefaciens*, the 480 bp long 16S rRNA region (V3-V4 region) contained 12 variable base sites (Fig. 4A and Fig. S3A), which were detected in only 4 of 116 genomes of this species (Fig. 4A), and the SNP frequency at variable sites within the 4 genomes ranged from 0.21%-1.46% (Fig. S3C red column). In contrast, the 500-nucleotide *gyrA* region (positions 350-850) contained 59 variable sites (Fig. 4B and Fig. S3B). The variable sites were detected in 109 of 116 genomes (Fig. 4B), and the frequency of SNPs at the variable sites ranged from 0.4% to 6.4% (Fig. S3C blue column).

**Figure 4.**
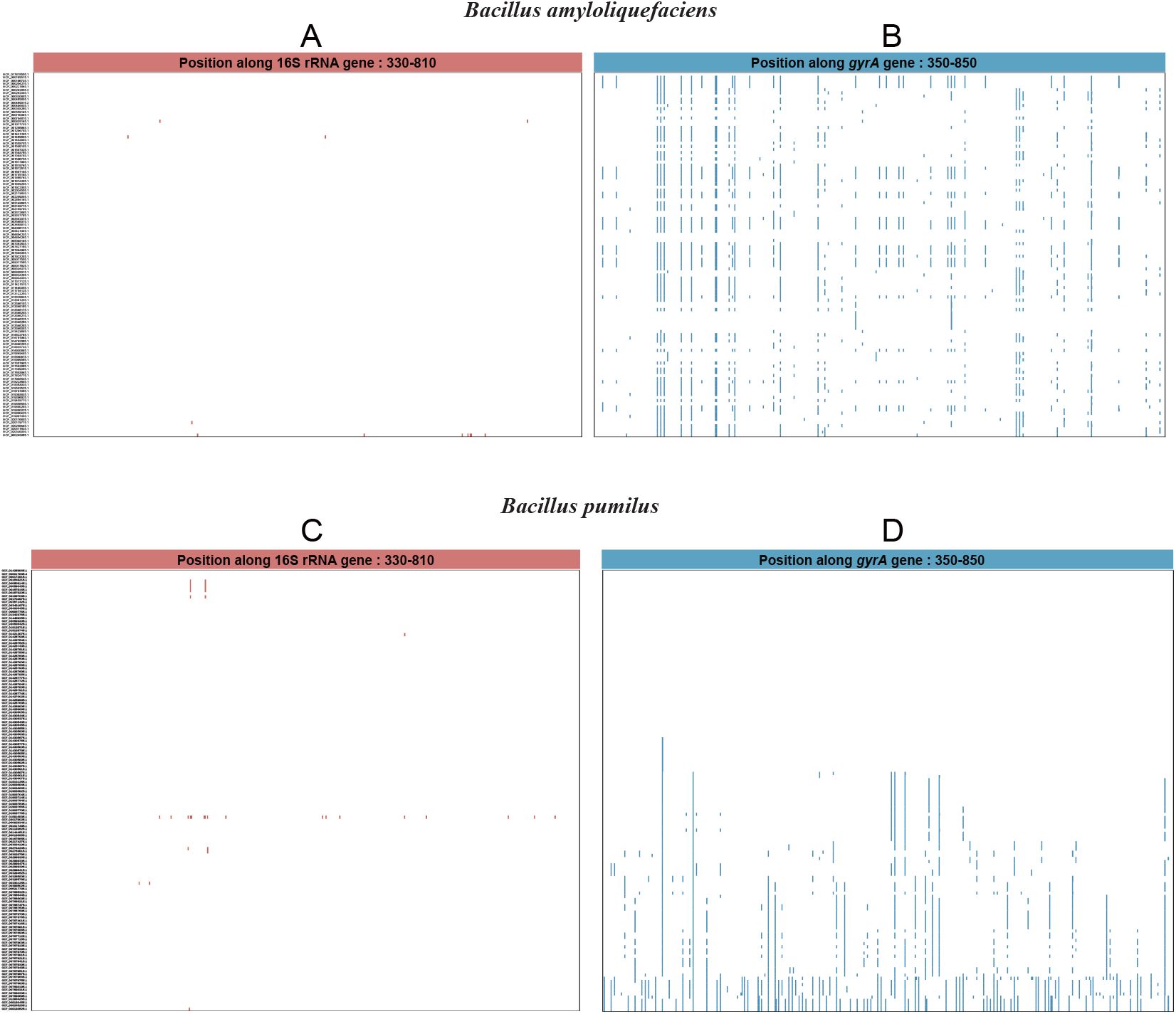
Alignment of the 16S rRNA and *gyrA* sequences of *B. amyloliquefaciens* (116 genomes) and *B. pumilus* (140 genomes). Sequences were aligned using L-INS-I method of MAFFT (v7.487). Analysis involved positions along 16S rRNA from 330-810 and along *gyrA* from 350-850. On the left side of the graph GCF reference numbers of genomes were displayed, with reference sequence GCF_017815555.1 displayed at the top for B. *amyloliquefaciens* and GCF_014269965.1 for *B. pumilus*. The display of base site variation was drawn using MEGA (v.5.05). The colored line (red or blue) indicates a variation of specific base sites as compared to the reference sequence.

In *B. pumilus*, alignment of the 16S rRNA V3-V4 region revealed 21 variable base sites (Fig. 4C and Fig. S4A), but again only in 11 out of 140 genomes (Fig. 4C). The frequency of SNPs at variable sites ranged from 0.21% to 3.54% (Fig. S4C red column). In contrast, in the *gyrA* gene 130 of the base sites were variable (Fig. 4D and Fig. S4B) and these were found in 87 of 140 genomes, with SNP frequencies at variable sites ranging from 0.2% to 15.8% (Fig. S4C blue column). However, in *B. pumilus* nearly 50% of the genomes examined had 100% identity in the *gyrA* gene, indicating high degree of relatedness between genomes that may require sequencing of additional marker genes for clonality verification and strain typing.

We also observed that the variation of the *gyrA* gene is quite different in different *Bacillus* species, which could be a bias of the NCBI database or a property of certain species. For example, *B. megaterium*, showed lower diversity with 3 bases of variation in the 16S rRNA alignment region (V3-V4 region) (Fig. S5A and Fig. S6A). Although 48 of 117 genomes showed polymorphism, the maximum SNP frequency at variable sites of *B. megaterium* genomes was only 0.42% (Fig. S6C red column). The *gyrA* gene was again more polymorphic with 58 bases of variation (Fig. S5B and Fig. S6B) occurring in 99 of 117 genomes, with SNP’s frequencies at variable sites ranging from 0.2% to 2.8% (Fig. S6C blue column).

The variation divergence between the two genes of *Bacillus anthracis* was much lower than in the three *Bacillus* species described above. Although we identified 13 variable base sites in the 16S rRNA V3-V4 region and 65 variable base sites in the *gyrA* gene region (Fig. S5C,D and Fig. S7A,B), these single nucleotide polymorphisms (SNPs) occurred in only 7 and 18 of 226 genomes, respectively. Moreover, the SNP frequencies at variable sites in 7 and 18 of *B. anthracis* genomes were also low: 0.21%-1.04% and 0.2%-7.4%, respectively (Fig. S7C).

Overall, our results showed that within *Bacillus* species the frequency of SNPs in the *gyrA* gene was consistently much higher than in the 16S rRNA (Fig. 4 and Fig. S5). We therefore suggest that the *gyrA* gene provides better resolution than 16S rRNA for identification and typing of *Bacillus* isolates at the subspecies level. This is particularly true for *B. amyloliquefaciens, B. pumilus* and *B. megaterium* but less so for *B*. anthracis (Table 1).

**Table 1.**
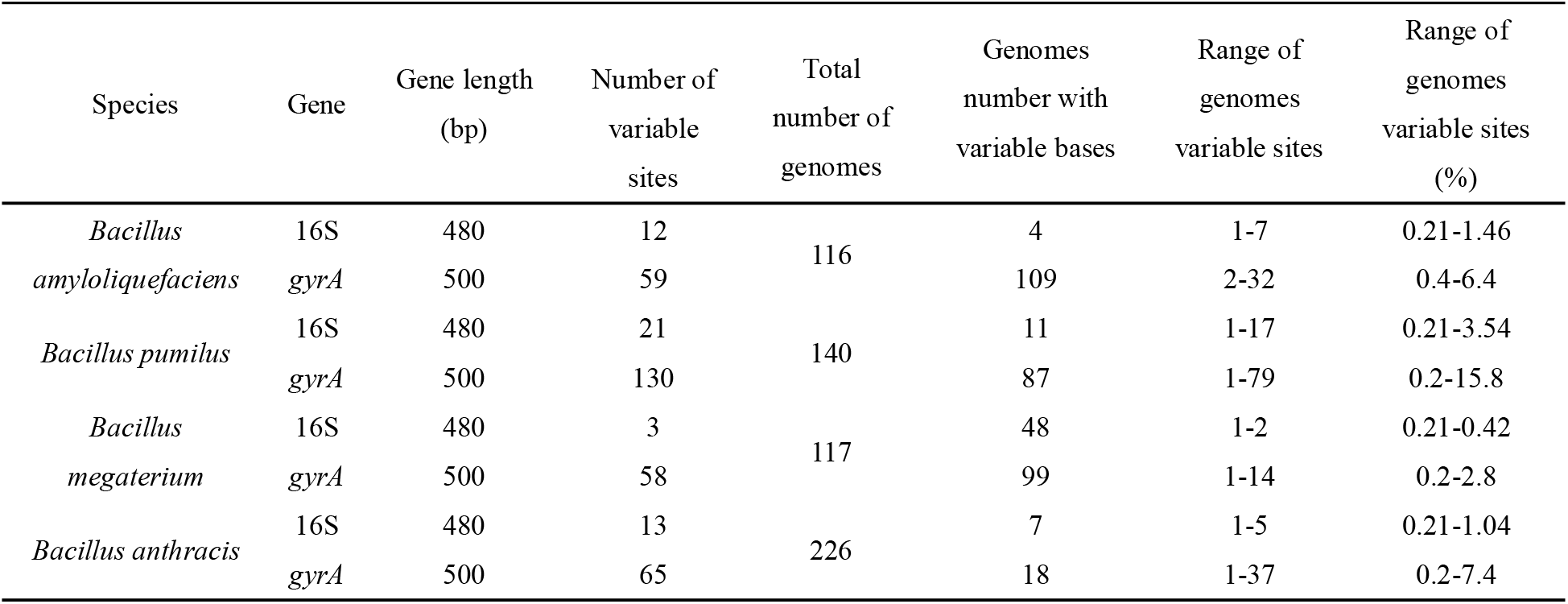
Polymorphisms of the 16S rRNA and *gyrA* gene.

### The resolution power of *Bacillus* mock community *gyrA* amplicon sequencing

Our results above suggest that the primer pair *gyrA*3 could be used as a molecular marker for diversity analysis of *Bacillus*. Next, we aimed to design a mock community to test the efficacy of the *gyrA*3 primer and used the general 16S rRNA primers (V3-V4) as a positive control. For better comparison, we designed two additional primer pairs, *gyrA*4 and *gyrA*5, which are very close to the position of the *gyrA*3 in the *gyrA* gene (Fig. S8A). We selected eight strains belonging to four species (*B. altitudinis, B. licheniformis, B. velezensis* and *Lysinbacillus pakistanensis*) and successfully amplified their *gyrA* gene by *gyrA*3, *gyrA*4 and *gyrA*5 primer pairs in a routine PCR for selected strains (Fig. 5A). Next, we artificially mixed eight indicated strains in equal proportions to obtain a mock community and then retested the resolution power of the three *gyrA* primer pairs and the 16S rRNA-specific primers. Our goal was to test whether the primer pair is suitable for determining the diversity of mock community (Fig. 5). Sequencing of the 16S rRNA amplicons showed that 16S rRNA primers could distinguish only five units, as strains LY1 and LY18, LY37 and LY43, and LY39 and LY48 had identical V3-V4 nucleotide sequence (Fig. 5B). The *gyrA* primer pairs were capable of resolving six units, but *gyrA*4 and *gyrA*5 produced amplicons of variable abundance and preferentially amplified LY35 and LY2, respectively(Fig. 5C-E). In comparison, primers *gyrA*3 also amplified six fragments but the relative abundance of these amplicons was more uniform (Table S6) with the exception of LY2 strain (Fig. 5C). Our data suggest that *gyrA*3 has potential for Illumina amplicon sequencing of more complex *Bacillus* communities.

**Figure 5.**
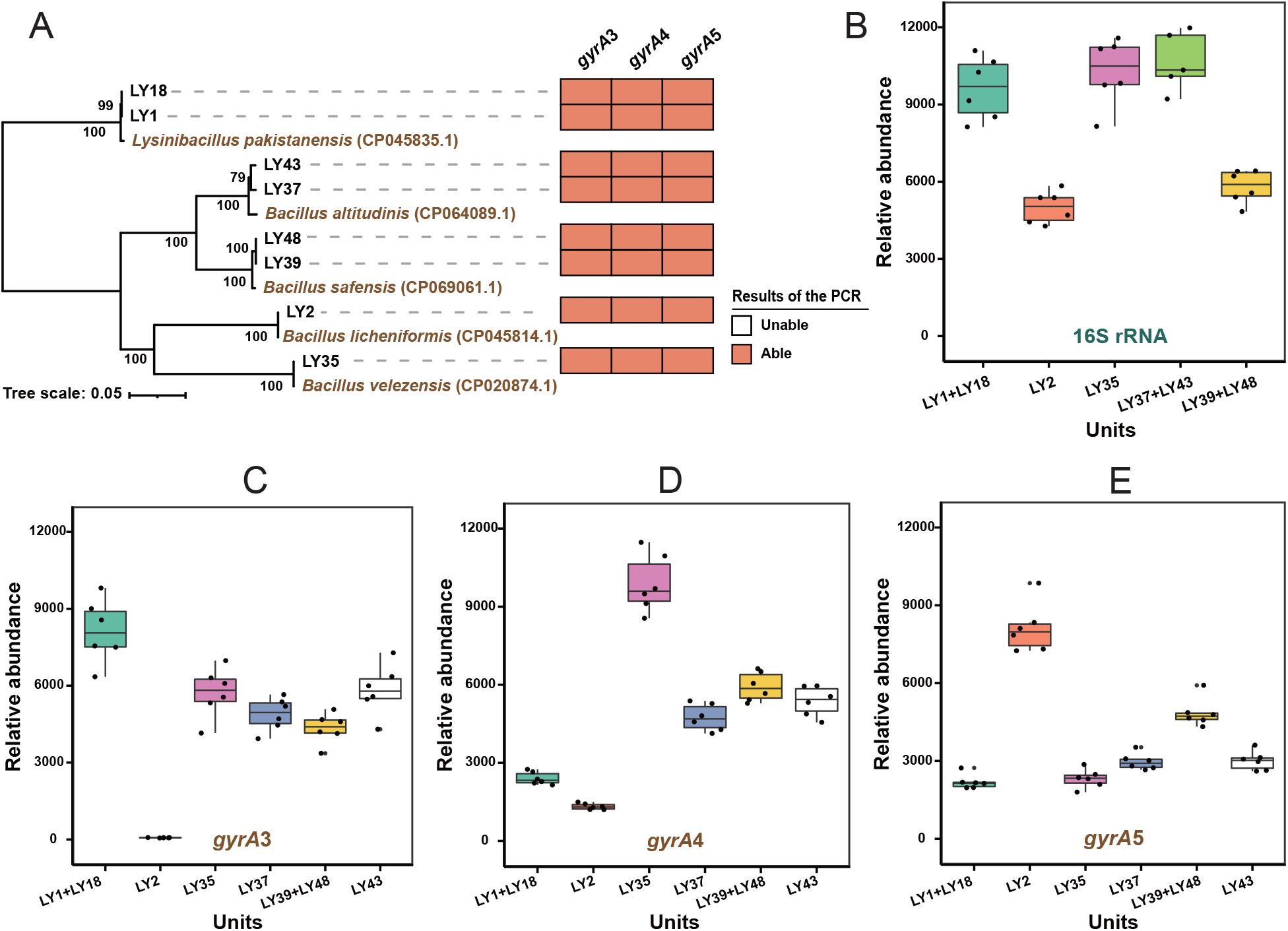
The results of amplicon sequencing for the mock community including eight *Bacillus* strains. (**A)** PCR amplification of eight *Bacillus* strains by *gyrA* gene primer pairs. The orange square indicates positive and white square unsuccessful amplification. The amplicon sequencing results of 16S rRNA gene (**B)** and *gyrA* gene using primer pairs *gyrA*3 (**C)**, *gyrA*4 (**D)**, and *gyrA*5 (**E)** for the mock community. The relative abundance represented by reads numbers for each unit as indicated on X axis. The phylogenetic tree was contracted based on the complete *gyrA* genes (2469bp) (Dataset S3) by using Neighbor-Joining method in MEGA 5.05 software. The reliability of clades was tested by the 1000 bootstrap replications. The box plots were drawn by ggplot2 R package (v.3.2.1).

## Discussion

It is believed that the diversity of microorganisms in nature is immense, and therefore its detection still remains a challenge (Widder et al., 2016). Molecular tools (e.g., for specific amplification of marker genes) combined with high-throughput sequencing are expected to open the door to the vast diversity of microorganisms (Klindworth et al., 2013). Here, we systematically investigated the potential of the *gyrA* gene as a marker gene for taxonomic typing of *Bacillus* isolates and assessment of *Bacillus* community composition by amplicon sequencing. The novel *gyrA*3 primer pair is capable of detecting 76 *Bacillus* species by virtual and colony PCR; hence, our results suggest that the *gyrA* gene is a good phylogenetic marker for detecting intra- and interspecific diversity of the genus *Bacillus*.

Sequencing of 16S rRNA amplicons on the Illumina platform has been the mainstream method of studying microbial communities, but this gene also shows shortcomings, especially when we aim to determine diversity within a genus or species (Amir et al., 2017; Callahan, McMurdie, & Holmes, 2017; Callahan et al., 2016; R. Edgar, 2016a, 2016b; R. C. Edgar, 2013). This is particularly true for *Bacillus* species, which exhibit very low interspecific variability (Vos et al., 2012). Although 16S rRNA is widely used as a molecular marker for bacterial community analyses (Clarridge, 2004), its amplicon sequencing can only describe community diversity at the genus level (Gupta et al., 2019). In contrast, faster evolution and consequently higher diversity of the *gyrA* gene (Timmis & Ramos, 2021) suggests that this gene might provide higher phylogenetic resolution than the 16S rRNA gene within the genus *Bacillus*.

In our study, the differential effect of the *gyrA* gene on *Bacillus* interspecies and several *Bacillus* species was better than that 16S rRNA (Fig. 1). Moreover, the results of comparative sequence analysis (including the 16S rRNA and the *gyrA* gene regions) of the three species: *B. amyloliquefaciens, B. pumilus*, and *B. megaterium*, support this prediction, as we find wider range of SNPs in the *gyrA* genes than in the 16S rRNA (Fig. 4, Fig. S5 and Table 1). Our results are consistent with findings that housekeeping genes, including *gyrA*, evolve much faster than 16S rRNA genes and are suitable for the identification and typing of closely related species (Poirier et al., 2018) and for the intraspecific resolution of isolates, as previously shown for *B. subtilis* (Ansaldi et al., 2002; Roberts et al., 1994).

To date, there have been only a few reports in which conserved genes (e.g. *rpoB* and *gyrB*) have been used as templates for amplicon sequencing of microbial communities (Ogier et al., 2019; Poirier et al., 2018; Vos et al., 2012). These reports show that sequencing of conserved protein coding genes provides a more accurate description of bacterial community composition than 16S rRNA sequencing. Specifically, the *rpoB* gene has been used in addition to the 16S rRNA molecular marker for high-throughput sequencing studies of species diversity in the *Proteobacteria* phylum (Vos et al., 2012).

Although it remains to be investigated whether *gyrA* is suitable as a molecular marker for the majority of *Bacillus* species, we identified here its resolution power for 76 *Bacillus* species (Fig. 3) and propose that the diversity of the *Bacillus* community can be more accurately assessed by combining 16S rRNA and *gyrA* amplicon sequencing. As a molecular marker, the housekeeping gene *gyrA* also has some limitations. Due to the high variability of *gyrA* gene sequences, it was difficult to design universal primers that can amplify the entire *Bacillus* genus, as has been previously described for the *Escherichia* genus (Johnning, Kristiansson, Fick, Weijdegård, & Larsson, 2015). Although *gyrA*3 significantly improved the amplification efficiency of *Bacillus* species compared to previous primers, not all *Bacillus* species could be detected (Fig. S2 and Fig. 3). On the other hand, the *gyrA* amplicon region we selected was well suited for resolving subspecies of *B. amyloliquefaciens, B. pumilus*, and *B. megaterium*. For *B. anthracis*, which is known for its low diversity (Lista et al., 2006) intraspecific resolution was limited (Fig. 4 and Fig. S5). Using amplicon sequencing, *gyrA*3 also had its own blind area in the detection, for example, it was not good at distinguishing between LY1 and LY18, LY39 and LY48 (Fig. 5). This could be due to the extremely high similarity of the *gyrA* genes in selected genomes, some of which have even identical *gyrA* sequences. However, the *gyrA*3 distinguished very well two strains with highly similar *gyrA* genes such as LY37 and LY43, which 16S rRNA did not (Fig. 5). Although primers for amplicon sequencing of the *gyrA* gene had certain limitations, such as not amplifying all selected targets or not reflecting the abundance of added DNA, this study has put forward the advantages that *gyrA*3 covers the broadest diversity of *Bacillus* species reported to date.

In summary, this study investigated the application of the *gyrA* gene as a molecular marker in *Bacillus* subspecies typing and high-throughput sequencing of the *Bacillus* mock community. The greater ability of *gyrA*-based analyses to distinguish *Bacillus* strains at the subspecies level should increase resolution and provide more reliable results for the ecological studies of the genus *Bacillus*. We believe that the primer pair will have broad applications in the *Bacillus* research.

### Materials and Methods Strains and culture condition

The 127 strains used in the study were strains of *Bacillus* and related genera of *Bacillus*, most of which were isolated from soil. Some strains labelled with ACCC are deposited in the Agricultural Culture Collection of China (http://www.accc.org.cn) (Table S4).

All strains were grown at 30 L in low-salt Luria-Bertani medium (LB), containing 10 g tryptone, 5 g yeast extract, and 3 g NaCl per litre.

### Nucleotide diversity (Pi) analysis

For interspecies analysis, the whole sequences of the 16S rRNA and the *gyrA* genes of 20 *Bacillus* species (361 genomes) were downloaded from the NCBI genome database (Table S1). For intraspecies analysis, three species were selected and whole sequences of 16S rRNA and *gyrA* gene were downloaded from the database: *Bacillus amyloliquefaciens* (88 genomes), *Bacillus licheniformis* (71 genomes) and *Bacillus pumilus* (97 genomes) (Table S1). Alignment of these sequences was conducted using online alignment tool Kalign on EMBL (https://www.ebi.ac.uk/Tools/msa/). Subsequently, nucleotide diversity (Pi) was estimated with DnaSP 6 (v.6) using a window size of 100 bp and a step size of 10 bp (Rozas et al., 2017).

### DNA extraction of strains, PCR and gel electrophoresis

Genomic DNA was extracted using the Omega Bacterial DNA Kit D3350 (Omega, Bio-tek, Norcross, GA, USA), and the concentration and quality of DNA were determined using a NanoDrop 2000 spectrophotometer (Wilmington, DE, USA).

The reaction mixture for PCR amplification was prepared in 25 μL containing 1 μL of DNA, 2 μL of each primer (the primer pairs *gyrA*1 were *gyrA*1-F: 5′-CAGTCAGGAAATGCGTACGTCCTT-3’ (Roberts et al., 1994) and *gyrA*1-R: 5′-GTATCCGTTGTGCGTCAGAGTAAC-3’ (Ansaldi et al., 2002), the primer pairs *gyrA*2 were *gyrA*2-F: 5′-CAGTCAGGAAATGCGTACGTCCTT-3’ (Roberts et al., 1994) and *gyrA*2-R: 5′-CAAGGTAATGCTCCAGGCATTGCT-3’ (Roberts et al., 1994), the primer pairs *gyrA*3 were *gyrA*3-F: 5′-GCDGCHGCNATGCGTTAYAC-3’ and *gyrA*3-R: 5′-ACAAGMTCWGCKATTTTTTC-3’, the primers for the 16S rRNA gene were 27F: 5′-AGAGTTTGATCCTGGCTCAG-3’ and 1492R: 5′-GGTTACCTTGTTACGACTT-3’), 12.5 μL Green Taq Mix (http://www.vazyme.com), and 7.5 μL deionized water.

PCR was performed under the following conditions: Predenaturation at 94 □ for 5 min; denaturation at 94 □ for 30 s; annealing at 50 □ for 30 s; elongation at 72 □ for 40 s (35 cycles); and elongation at 72 □ for 7 min.

### In Silico PCR

The *gyrA* sequence database was constructed as a virtual community, containing 5062 full-length sequences from the National Center for Biotechnology Information (NCBI) belonging to 226 *Bacillus* species (Table S2) (the method for obtaining the database sequences is described in detail in the SNP analysis section below). First, the degenerate primer pair *gyrA*3 was converted to primers that do not contain degenerate bases, and the converted *gyrA*3 is listed in Table S7. Subsequently, the primer pairs *gyrA*1, *gyrA*2 and the converted *gyrA*3 were aligned with the *gyrA* database using NCBI-blast+ software (v.2.9.0). The match at 18 bases of both, the forward and reverse primer, was considered amplifiable by the primer pair.

### Phylogenetic analysis

In this study, phylogenetic analysis of genes was performed using MEGA (v.5.05) (Tamura et al., 2011) for Neighbor-Joining and the reliability of clades was tested using 1000 bootstrap replicates. Furthermore, annotation and beautification of trees was performed using programs available at the iTol online site (https://itol.embl.de) (Tamura et al., 2011).

### SNP analysis

A total of 734 available genomes of 4 different *Bacillus* species were downloaded from the NCBI database using the NCBI-genome-download script (https://github.com/kblin/ncbigenome-download/) (Table S5) including 124 genomes of *B. amyloliquefaciens*, 167 genomes of *B. pumilus*, 150 genomes of *B. megaterium* and 293 genomes of *B. anthracis*. Then, the location information of 16S rRNA and *gyrA* gene sequences on the genomes were determined by aligning 16S rRNA and *gyrA* sequences of the same species using NCBI-blast+ (v.2.9.0) and the 16S rRNA / *gyrA* gene sequences were extracted using the Fasta Extract tool in TBtools (v1.0986853) (Chen et al., 2020). After filtration, genomes that did not carry complete 16S rRNA and *gyrA* genes, were removed and 600 genomes were retained, including 116 genomes of *B. amyloliquefaciens*, 140 genomes of *B. pumilus*, 117 genomes of *B. megaterium* and 226 genomes of *B. anthracis* (Table S5).

The alignment of the 16S rRNA (base sites:330-810) and *gyrA* genes (base sites:350-850) within each species was performed using the L-INS-I method of MAFFT (v7.487) (https://mafft.cbrc.jp/alignment/software/). The two gene sequences (16S rRNA, *gyrA*) on the same genome were selected as representative sequences (the top sequence of each figure was the representative sequence), and the base mismatch sites on other sequences were marked with color after comparison with the representative sequence. The genomes of the representative sequences in different species were GCF_017815555.1 (*B. amyloliquefaciens*), GCF_014269965.1 (*B. pumilus*), GCF_002577645.1 (*B. megaterium*) and GCF_000007845.1 (*B. anthracis*) (Fig. 4 and Fig. S5).

### Construction of the DNA-based *Bacillus* mock community, amplicon sequencing and data analysis

For the *Bacillus* mock community, we selected 8 strains with known genome sequences. Genomic DNA from 8 strains was extracted and its quality and quantity determined. Eight genomic DNAs were pooled in equal amounts after being diluted to approximately the same concentrations.

The hypervariable region V3-V4 of the 16S rRNA gene was amplified with the universal primers 338F: 5′-CCTACGGRRBGCASCAGKVRVGAAT-3’ and 806R: 5′-GGACTACNVGGGTWTCTAATCC-3′. The *gyrA* gene from 8 strains of the mock community was amplified with primer pairs *gyrA*3 (see above), *gyrA*4 (F:5′-TAYGCRATGAGYRTHATYGT-3’ and R: 5′-TTBGTNGCCATHCCDACMGC-3’), and *gyrA*5 (F:5′-GCDGCNGCVATGCGTTAYAC-3’ and R: 5′-CGNAGRTYBGTAATDCCDTC-3’). Sequencing was performed on an Illumina Miseq PE300 instrument.

Raw data were processed using the UPARSE pipeline (http://drive5.com/usearch/manual/uparse_pipeline.html) (R. C. Edgar, 2013) to obtain the ZOTUs table. This included truncating and assembling, removing double end primers, filtering low-quality sequences, finding unique sequences, denoising, and creating the ZOTUs with 100% similarity, which were then used to create the ZOTUs table. Denoising (error-correction) of the amplicon reads was performed by using the Unoise3 algorithm (R. Edgar, 2016b) to identify and correct reads with sequencing errors and remove chimeras.

The eight strains of the mock community with known complete 16S rRNA and *gyrA* gene sequences were used to annotate the ZOTUs (Dataset S2 and S3). The annotation of ZOTUs sequences was performed with NCBI-blast+ (v.2.9.0). The ZOTUs of the same species were pooled into a single unit after annotation. Finally, we used box plots to show the community structure and the characteristics of the different primer pairs during sequencing. The box plots were drawn using R (v.4.0.3).

## Acknowledgements

This work was financially supported by the National Nature Science Foundation of China (31972512 and 42090064), the Fundamental Research Funds for the Central Universities (KYXK202009). The Innovative Research Team Development Plan of the Ministry of Education of China (Grant No. IRT_17R56). PS and IMM were supported by the Program Grant P4-0116 funded by the Slovenian national research agency (ARRS).

## Author contributions

YL and ZX designed the study; YL performed the experiments. YL, YM and PS analysed the data and created the figures. YL, PS and ZX wrote the first draft of the paper; PS, ZX, RZ, QS and IM revised the paper.

## Figure legends

**Figure S1** The agarose gel electrophoresis of DNA amplification products of three *gyrA* gene primer pairs: *gyrA*1 **(A)**, *gyrA*2 **(B)** and *gyrA*3 **(C)**. 1-7 indicates strains LY18, LY25, LY37, FZB42, SXL408, SXL277 and ACCC01043.

**Figure S2** Alignment of the 16S rRNA and *gyrA* sequences of *B. megaterium* (117 genomes) and *B. anthracis* (226 genomes). Sequences were aligned using L-INS-I method of MAFFT (v7.487). Analysis involved positions along 16S rRNA from 330-810 and along *gyrA* from 350-850. On the left side of the graph GCF reference numbers of genomes were displayed, with reference sequence GCF_002577645.1 displayed at the top for *B. megaterium* and GCF_000007845.1 for *B. anthracis*. The display of base site variation was drawn using MEGA (v.5.05). The colored line (red or blue) indicates a variation of specific base sites as compared to the reference sequence.

**Figure S3** Polymorphisms in the 16S rRNA and *gyrA* gene’s regions of *B. amyloliquefaciens* (116 genomes). **(A)** The proportion of variation at different base sites along 16S rRNA gene V3-V4 region. **(B)** The proportion of variation at the different base positions of the *gyrA* gene region. **(C)** The proportion of variants along 16S rRNA (red column) and *gyrA* gene (blue column) region in different genomes.

**Figure S4** Polymorphisms in the 16S rRNA and *gyrA* gene’s region of *B. pumilus* (140 genomes). **(A)** The proportion of variation at different base sites along the 16S rRNA gene **(B)** and the *gyrA* gene. **(C)** The proportion of variable base sites in 16S rRNA (red column) and *gyrA* (blue column) sequences in different genomes.

**Figure S5** Polymorphisms in the 16S rRNA and *gyrA* gene’s regions of *B. megaterium* (117 genomes). **(A)** The proportion of variation at different base sites along 16S rRNA gene V3-V4 region. **(B)** The proportion of variation at the different base positions of the *gyrA* gene region. **(C)** The proportion of variants along 16S rRNA (red column) and *gyrA* gene (blue column) region in different genomes.

**Figure S6** Polymorphisms in the 16S rRNA and *gyrA* gene’s region of *B. anthracis* (226 genomes). **(A)** The proportion of variation at different base sites along the 16S rRNA gene **(B)** and the *gyrA* gene. **(C)** The proportion of variable base sites in 16S rRNA (red column) and *gyrA* (blue column) sequences in different genomes.

**Figure S7** Description of *gyrA* gene primer pairs for amplicon sequencing. (**A)** The position of *gyrA* amplicons obtained by primer pairs *gyrA*3, *gyrA*4 and *gyrA*5. (**B)** PCR amplification of eight *Bacillus* strains by *gyrA* gene primer pairs. The orange square indicates positive and white square unsuccessful amplification. The phylogenetic tree was contracted based on the complete *gyrA* genes (2469bp) (Dataset S3) by using Neighbor-Joining method in MEGA 5.05 software. The reliability of clades was tested by the 1000 bootstrap replications.

