## Supplementary figures and images for "Housekeeping gene *gyrA*, a potential molecular marker for *Bacillus* ecology study"

### Figure S1

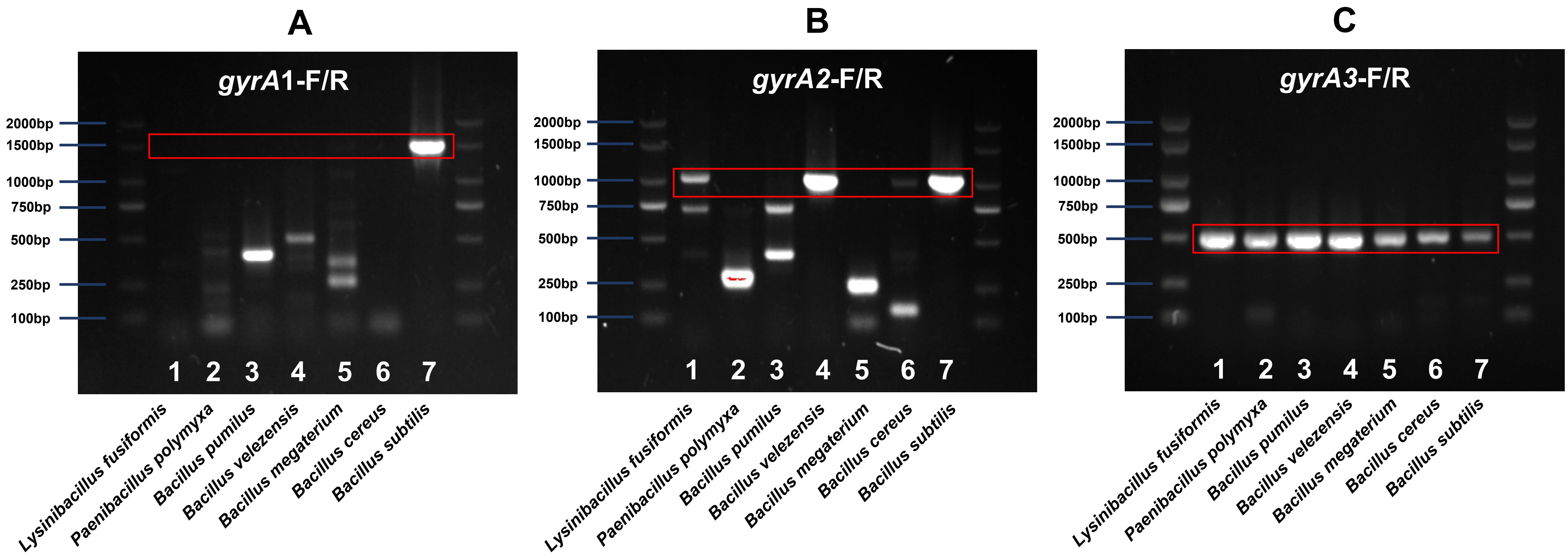

### Figure S2

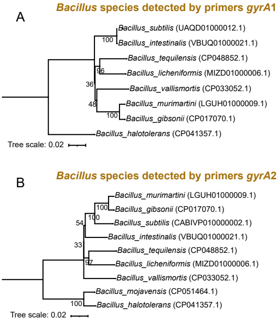

### Figure S3

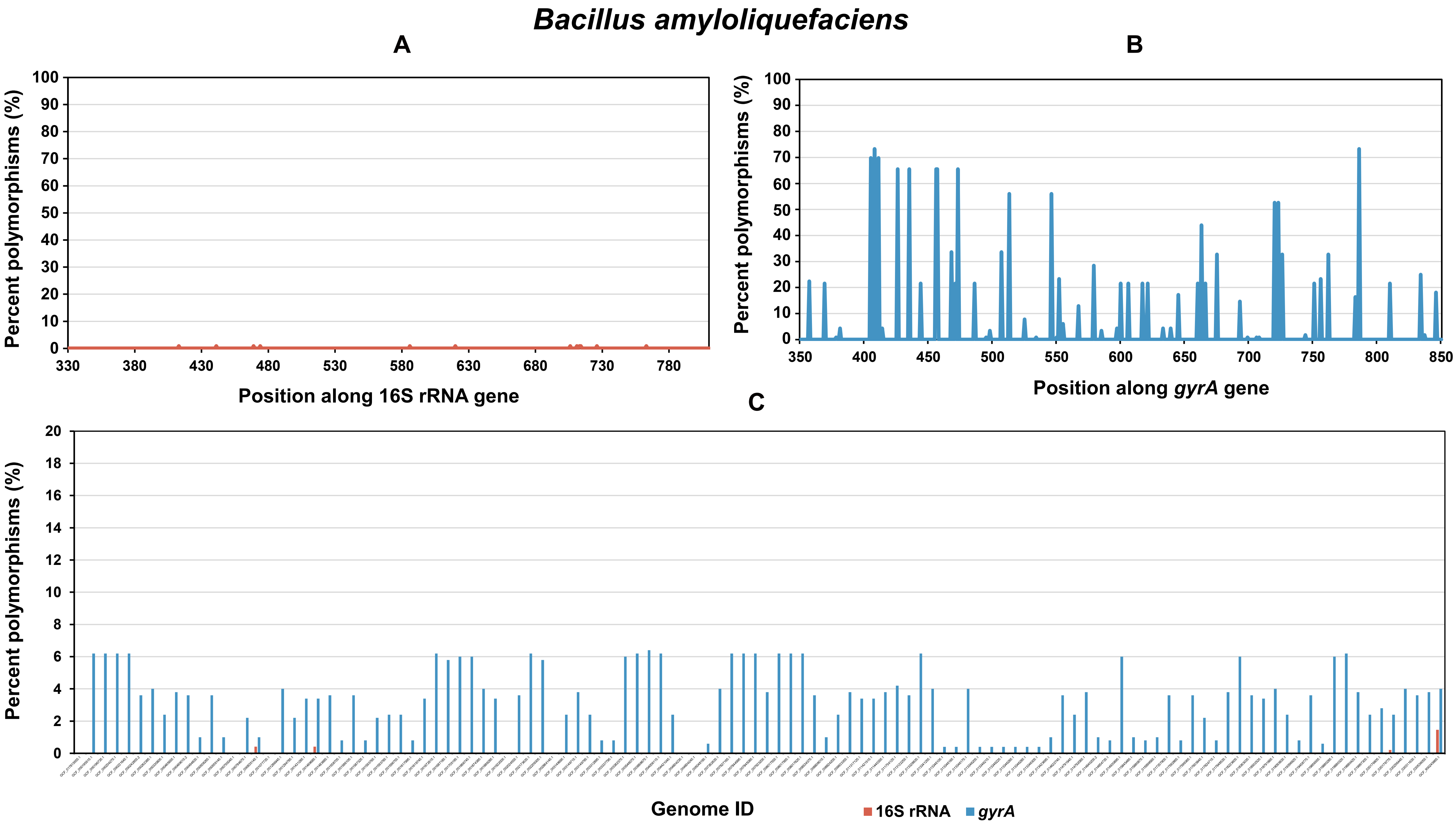

### Figure S4

#### *Bacillus pumilus*

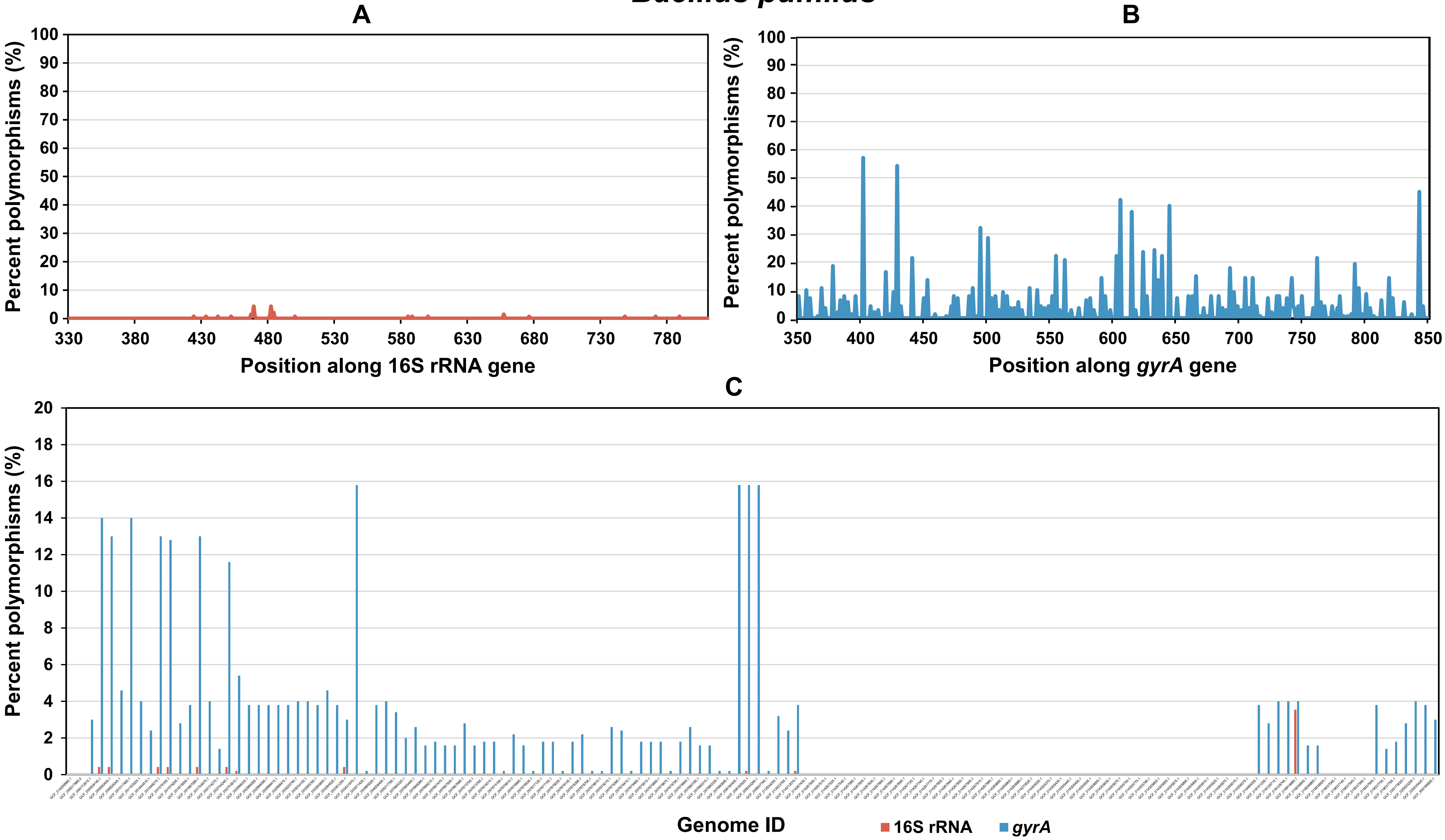

### Figure S5

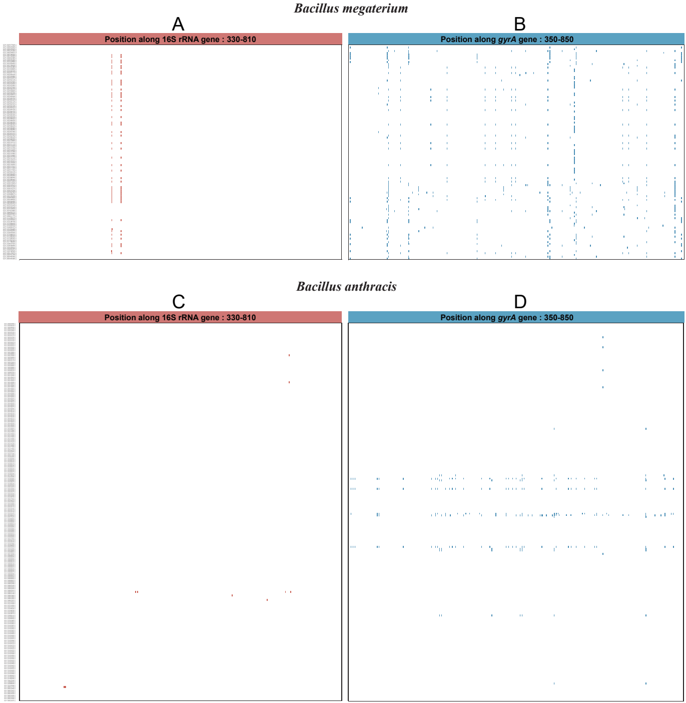

### Figure S6

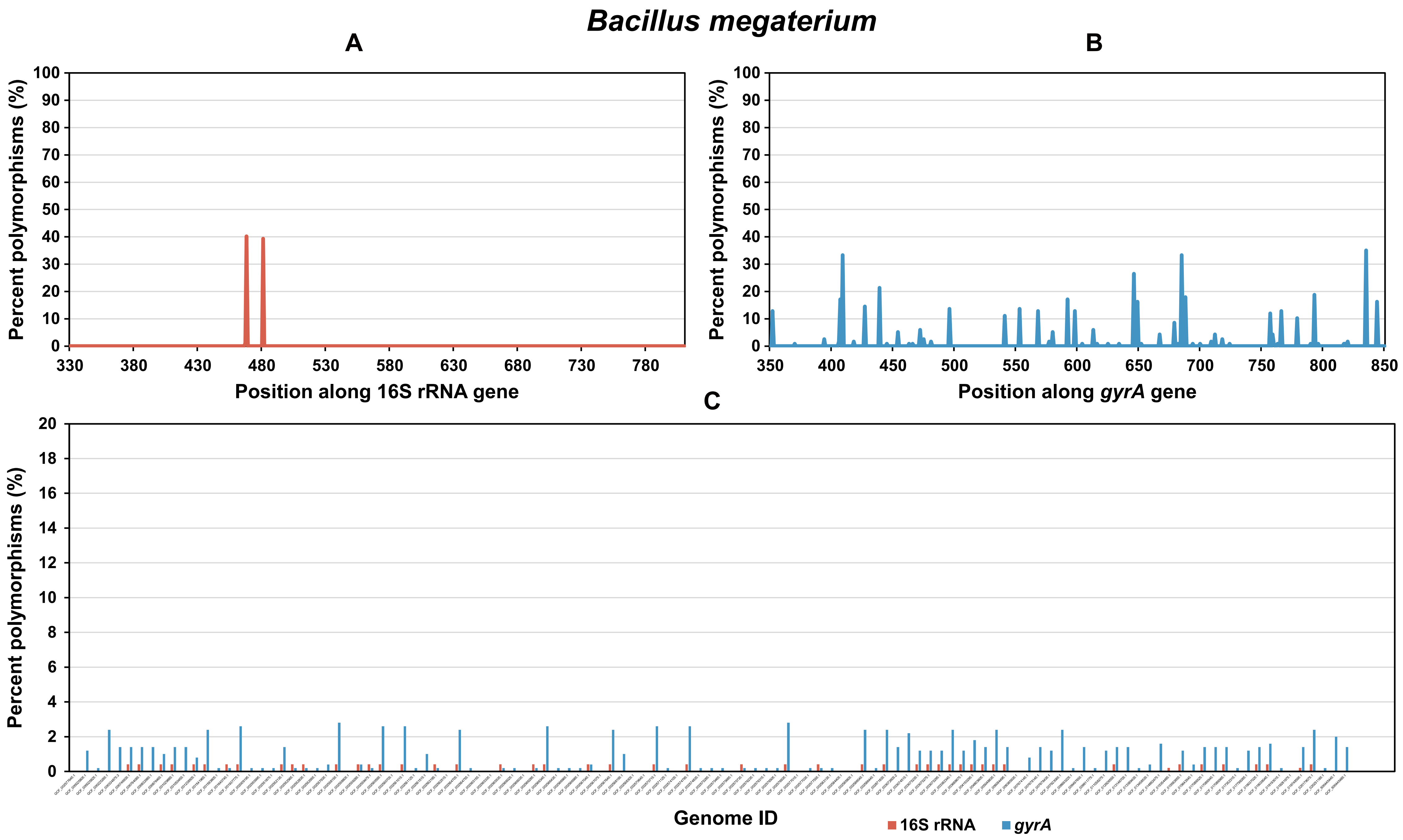

### Figure S7

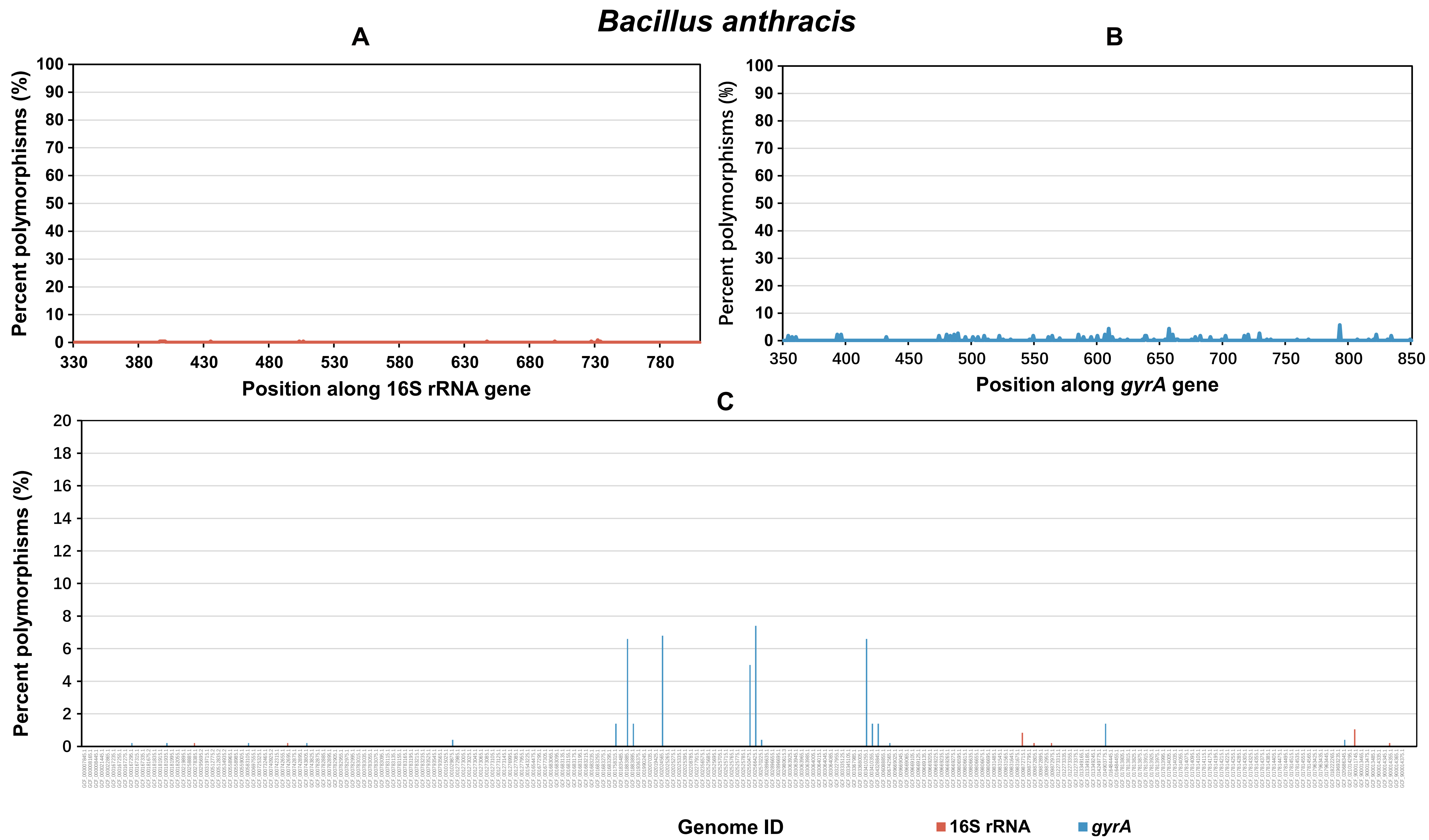

### Figure S8

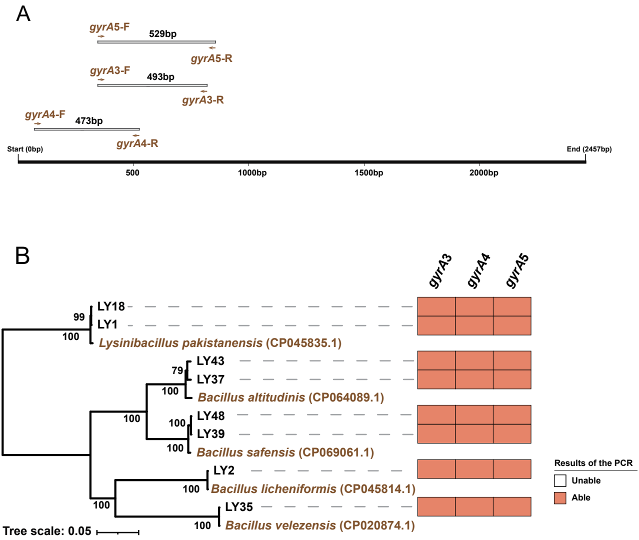
